# Towards Staining Independent Segmentation of Glomerulus from Histopathological Images of Kidney

**DOI:** 10.1101/821181

**Authors:** Robin Liu, Lu Wang, Jim He, Wenfang Chen

## Abstract

This paper introduces a detection-based framework to segment glomeruli from digital scanning image of light microscopic slide of renal biopsy specimens. The proposed method aims to better use the precise localization ability of Faster R-CNN and powerful segmentation ability of U-Net. We use a detector to localize the glomeruli from whole slide image to make the segmentation only focus on the most relevant area of the image. We explored the effectiveness of the network depth on its localization and segmentation ability in glomerular classification, and then propose to use the classification network with enhanced ability of localization and segmentation to construct and initialize a segmentation network. We also propose a weakly supervised training strategy to train the segmentation network by taking advantage of the unique morphology of the glomerulus. Both strong initialization and weakly supervised training are used to resolve the problem of insufficient and inaccurate data annotations and enhance the adaptability of the segmentation network. Experimental results demonstrate that the proposed framework is effective and robust.

## 1 Introduction

Histological staining is commonly used in diagnostic renal pathology, where biopsy specimens is stained to highlight important features of the tissue, i.e. Periodic acid-Schiff (PAS) staining is used to highlight basement membranes, while Masson’s trichrome (MT) staining is used to discriminate collagen fibers from muscular tissues. In some cases, differential staining, double staining or multiple staining are also used. Due to the variations in staining methods, staining intensity, thickness of the biopsy tissue section, slide scanners, and laboratories, significant differences of glomerular color, shape and texture exist among different whole slide images (WSIs), as shown in Fig. 1. In addition, the doctor’s operation also affects the shape of glomerulus, there can exist tangential glomerulus, complete glomerulus and incomplete glomerulus in a WSI.

**Fig. 1.**
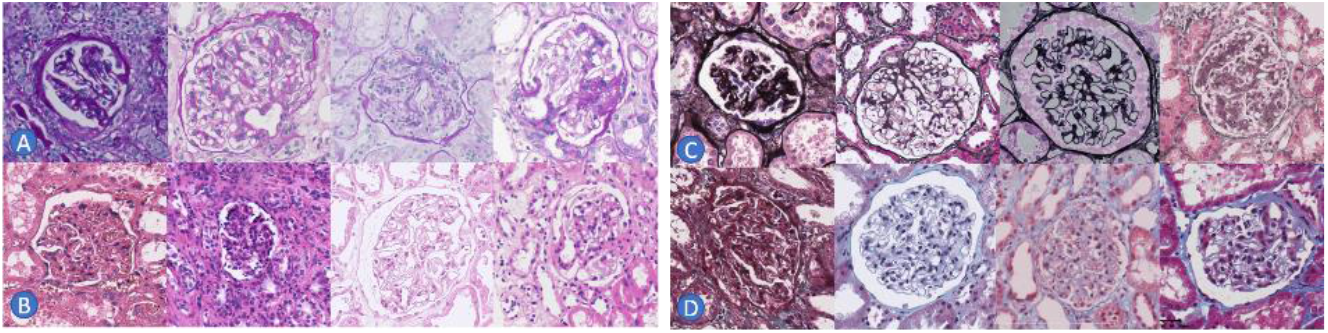
Stained glomeruli, A: Periodic acid-Schiff (PAS), B: Hematoxylin and eosin (H&E), C: Periodic acid-silver methenamine (PASM), D: Masson’s trichrome (MT)

More recently, several attempts [1, 2] have been discussed about the staining independent or staining invariant segmentation of glomeruli. The stain-translation solution [2] can only be applied to the glomeruli with well-defined boundary, and the target stain must be with similar texture to the source stain. As color, shape, and texture of the glomeruli differed greatly in various staining, for the first time, we propose a detection-based framework to segment the glomeruli from large WSI automatically. As shown in Fig.2, the framework consists of two steps in sequence: A Faster R-CNN [3] to localize and detect the glomeruli, and a convolutional neural network using modified U-shaped [4] architecture to segment the glomerulus-centered image. In the framework, several different ways are used to resolve the challenges we faced: 1) we use a detector to localize the glomeruli from large WSI to make the segmentation can only focus on the most relevant small area; 2) we use a sub-network with enhanced ability of localization and segmentation to construct and initialize the segmentation network; 3) we propose a weakly supervised training strategy to train the segmentation network by taking advantage of the unique morphology of the glomerulus.

## 2 Methods

### 2.1 Glomerular Detection

As a staining independent detector, it should be trained on the dataset which should cover as many kinds of staining as possible. Except several popular staining, we also generate several fake staining by transferring the color characteristics of the source staining to target staining [5]. Due to the limitation of computing resources and processing speed consideration, we down-sample all WSIs with 40x magnification to 5x magnification and the diameter of the glomeruli at this magnification is about 40 ∼ 150 pixels, which are larger enough to maintain the performance unchanged for detection. All the training patches with fixed-size of 800×800 pixels were generated by randomly cropping regions from the 5x WSIs, where the objects partially cropped at the edge of the region were marked as background and the patches with only background were also included in the training dataset. In Faster RCNN, the images are first fed into a region proposal network to get the proposed objects, then the followed classification and regression networks are used to refine them. The Inception-v2 [6] is used as the feature extractor of the Faster RCNN in our framework, which have enough receptive field to make the network can identify the very large glomeruli.

### 2.2 Segmentation Network

In order to resolve the issue of the limited availability of manually labeled glomeruli, we propose to train a network to perform glomerular classification using large scale image-level labels first, then use the pre-trained classification network as the encoder of the segmentation network. The pre-trained weights from the classification network will give the segmentation network a strong initialization in the training processes. VGG network [7] is used as the classification network in our framework, where the fully connected layers before the final classification layer are replaced by a global average pooling layer to enhance discriminative localization ability of the classification network [8].

Given amount of annotated bounding boxes of the glomeruli, the image-level labels including glomeruli and backgrounds are automatically generated from the WSIs at 10x resolution by cropping the patches to a fixed size 256×256, as shown in Fig. 2, where the background patches are randomly selected from the non-glomeruli area. Base on the generated large-scale training images, 4 VGG networks with different number of convolutional layers 8, 10, 13, and 16 were trained to investigate the effectiveness of the network depth on its localization and segmentation ability by using the class activation map [9]. By feeding 120 generated testing images to the 4 classification networks to obtain their activation maps, as shown in Fig. 3, the mean Dice score between the binarized activation maps and their ground truth mask was used to choose the network with most powerful localization and segmentation ability. Finally, the pre-trained classification network with 10 convolutional layers was selected as the encoder of the segmentation network.

**Fig. 2.**
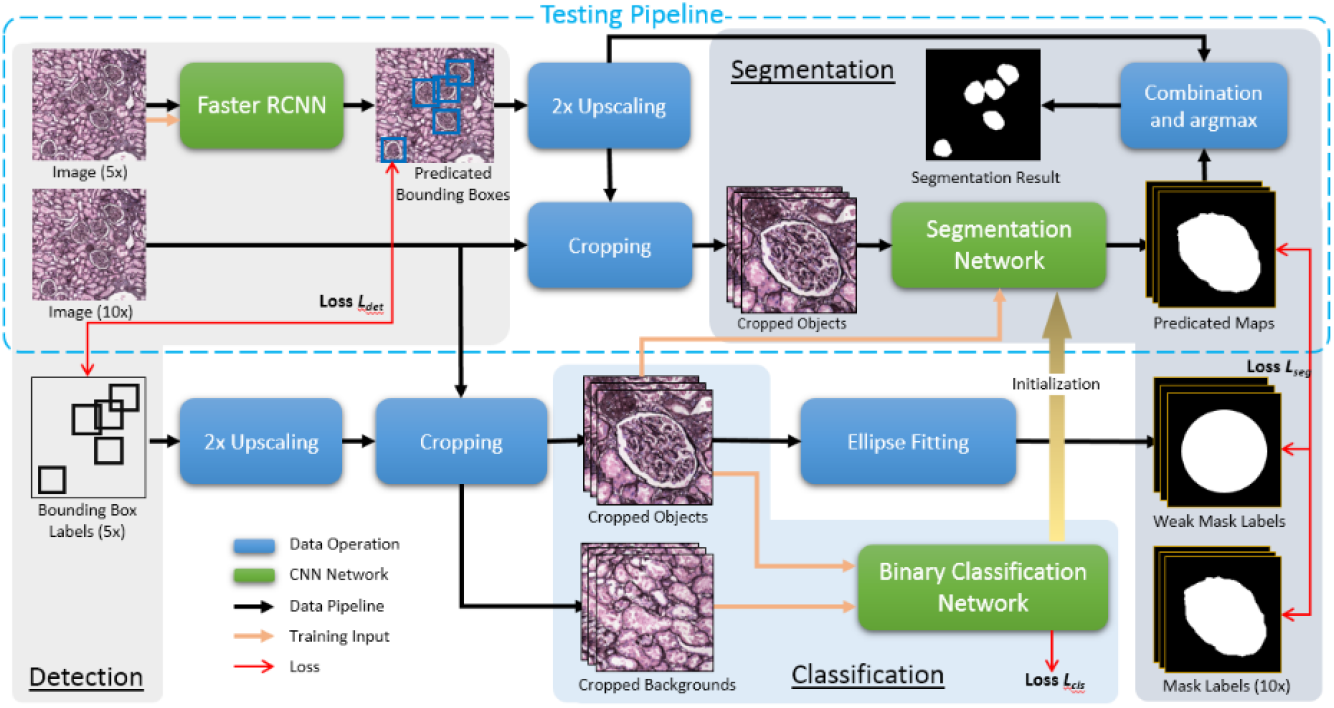
Detection based segmentation framework. The detection part outputs the bounding boxes and locations of each glomerulus, the following segmentation network segments each glomerulus on the glomerulus-centered image.

**Fig. 3.**
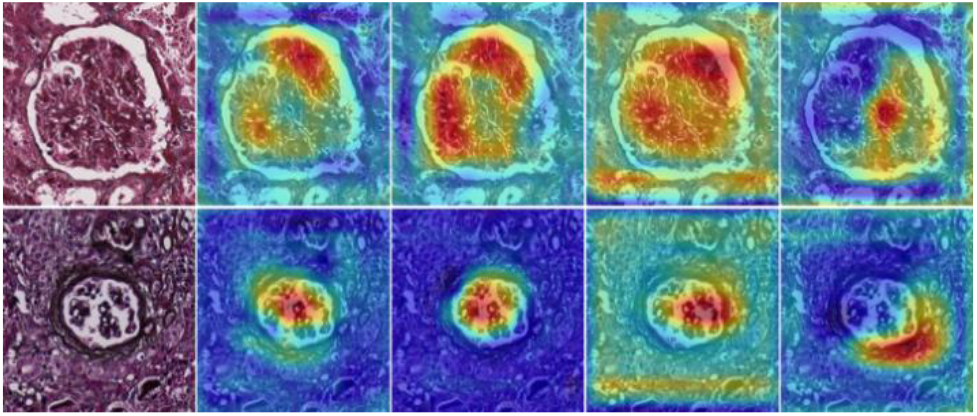
The figure shows the class activation maps across the networks with different number of convolutional layers 8, 10, 13, and 16 (from left to right) trained for glomerular classification.

Fig. 4 illustrates the architecture of the segmentation network, which starts with an encoder sub-network followed by a decoder sub-network and an output layer using sigmoid. Inspired by the iterative deep aggregation [10], aggregation nodes were inserted into the segmentation network to merge the features from shallow layer to deep layer and hence the short skip connection replaced the long skip connection. The aggregation of channels, scales, and resolutions will help to reduce the semantic gap between the features extracted from the encoder and decoder and enhance the representative capacity of the deep features to represent the consistent information among various stains. In our architecture, different number of aggregation nodes 4 and 10 are experimented to find which style of aggregation would achieve better performance.

**Fig. 4.**
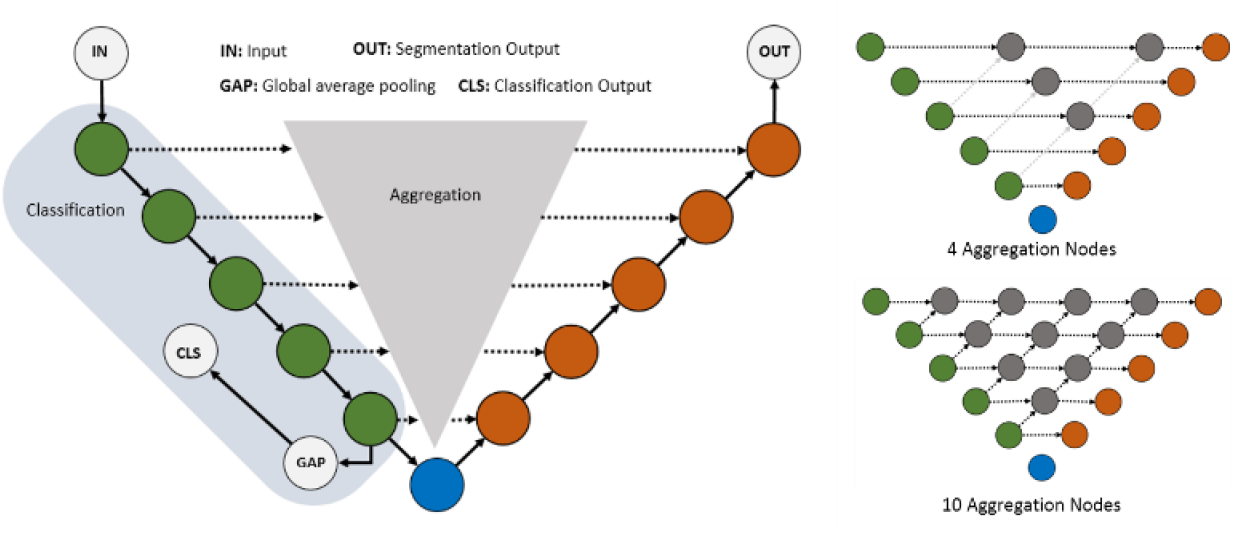
The architecture of the segmentation network. The input size: 256×256×3, the output size: 256×256×1, encoder block: conv-bn-relu + conv-bn-relu + max-pool, bottleneck: conv-relu + conv-relu + dropout, decoder block: up-conv-relu + concatenate + conv-relu + conv-relu, aggregation node: up-conv-relu + concatenate + conv-relu, where convolution (conv) uses 3×3 kernel size, up-convolution (up-conv) and max-pooling (max-pool) uses 2×2 kernel size, batch normalization (bn) is used in encoder block, rectified linear unit (relu) is used as activation function for all convolutions layers except the output layer, dropout ratio is 0.5.

### 2.3 Weakly Supervised Training Strategy

Since most of the glomeruli are egg-shaped, in this work we propose to use ellipses to fit the contour of annotated bounding boxes of glomeruli and use the fitting results as weak labels to train the segmentation network. Our goal is to train a segmentation network in a weakly supervised setting using a small fully supervised dataset 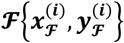 and a large weakly supervised dataset 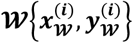, where ***x*** represents an image, ***y*** represents label of the image, ***i*** indicate the index of the image. The network with the strong initialized encoder is first trained on the small fully supervised dataset. Then it is fine-tuned iteratively by adding more and more weak labels into the training dataset.

Let suppose current training set is 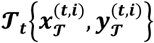, where ***t*** indicates the iteration index. In each iteration, a candidate set 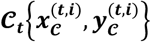 with same number of images of training set **𝒯** is chose from the weak dataset **𝒲** and fed to the network to obtain the predications 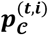, and then postprocessing including hole-filling and keeping largest object is applied to the predications. The Dice score 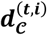 between the processed predications and their weak labels are calculated, and the new labels is determined by a threshold ***d***_***th***_,

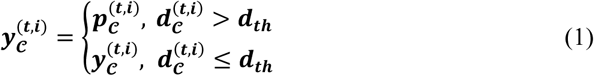

After the labels updated in the candidate set, the new training set for next iteration is created by combined the old training set with the candidate set.

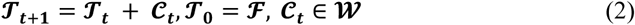

In the whole training procedure, the training / fine-tuning will iterate several times, and the number of iterations is determined by the size of the weak dataset.

## 3 Results and Discussion

### 3.1 Dataset and Training

#### Detection

Total 4056 glomeruli from 100 WSIs at 5x resolution were used to train the faster RCNN using SGD optimizer (with momentum of 0.9). The training dataset includes 4 kinds of stains H&E, PAS, PASM, and MT.

#### Classification

Total 4056 glomerulus-centered images and 4056 background images were used to train the classification network with binary cross-entropy as loss function.

#### Segmentation

Total 120 glomerulus-centered images with annotated mask and 1063 images with annotated bounding boxes were used to train the segmentation network. In the whole training procedure, the training / fine-tuning iterated 4 times with ***d***_***th***_ = **0. 8**. In each iteration, the network is trained / fine-tuned by optimizing the Dice loss function using Adam optimizer with a learning rate of 0.0001.

#### Experiments

Total 2145 glomeruli from 40 WSIs were used to evaluate the framework. The testing dataset also includes 4 kinds of stains (H&E, PAS, PASM, and MT). The color and texture of the glomeruli in the testing dataset are significantly different from the training dataset. The whole testing dataset was used to evaluate the detection, and the sliding window was applied to help to perform whole image detection. Total 405 glomeruli are selected to evaluate the segmentation. Except the original RGB images, their channel swapping and grayscale conversion were also used to evaluate the robustness of both detection and segmentation. In our experiments, 7 groups of images were totally used, includes the original RGB images, images swapped with any 2 color channels (i.e. RGB -> BGR), images swapped with all 3 channels (i.e. RGB -> GBR), images with single-channel (i.e. RGB -> G, RGB -> B, RGB -> R), and images mixed all 3 channels (i.e. RGB -> Grayscale). As the segmentation network only accepts images with 3 channels as input, we converted the single-channel image to 3 channels before feeding it to the network. Please be noted that we haven’t used any channels swapped or grayscale images in the training data.

### 3.2 Glomerular Detection

We measure the detection accuracy by precision, recall, F1-score at IoU (Intersection over Union) threshold 0.5 for each testing groups. Compared the results to the original RGB group (Table 1), there are almost no performance degradation observed in the testing groups which swapped 2 and 3 channels. It demonstrates that the deep features extracted by the detector represent the consistent information well among various colors or stains. Recall of the grayscale and single channel groups decreased but precision increased, which indicates that the features extracted by the deep network from the images with mixed channels or single channel are not strong enough to provide comprehensive information to the final classifier.

**Table 1.** Detection accuracy of different testing groups (IoU = 0.5).

|  | RGB | BGR | GBR | G | B | R | Gray | Avg. |
| --- | --- | --- | --- | --- | --- | --- | --- | --- |
| Precision | 0.965 | 0.965 | 0.959 | 0.983 | 0.988 | 0.990 | 0.989 | 0.977 |
| Recall | 0.971 | 0.968 | 0.956 | 0.894 | 0.914 | 0.897 | 0.903 | 0.929 |
| F1-score | 0.968 | 0.966 | 0.957 | 0.936 | 0.949 | 0.941 | 0.944 | 0.952 |

### 3.3 Glomerular Segmentation

For comparison, the U-Net [4], U-Net++ [11], and SE U-Net [12] were trained and fine-tuned using same loss function and the proposed weakly supervised training strategy. Table 2 lists the segmentation accuracy measured by mean Dice score of all the testing groups. Fig. 5 illustrates the ground truth together with results from U-Net, U-Net++, SE U-Net and proposed framework. We observed when there is no strong initialization applied, our architectures (Ux-A4 and Ux-A10) can’t perform better than other architectures in some testing groups (i.e. GBR, B, and R). But when the strong initialization applied, the performances of both our architectures (Ux-A4 /s and Ux-A10 /s) were significantly improved and they achieve a large performance gain over all other networks. The strong initialization, as is provided by the classification network trained from large dataset, make the segmentation network more easily to converge to the global minimum. As seen, the use of weakly supervised training strategy improved the Dice score for all segmentation networks including the U-Net, U-Net++, and SE U-Net. Specially for our architecture with 10 aggregation nodes (Ux-A10 /s/w), the strategy led very consistent performance across all testing groups. We also compared the performance of our architecture with different number of aggregation nodes. The results shown that the network with 10 aggregation nodes gives better performance than the one with 4 aggregation nodes.

**Table 2.** Mean Dice scores of different testing groups for U-Net, U-Net++, SE U-Net, and our architectures with 4 and 10 aggregation nodes (Ux-A4 and Ux-A10), where **/w** means network trained using weakly supervised training strategy, **/s** means strong initialization is applied to network before training.

| Method | RGB | BGR | GBR | G | B | R | Gray | Avg. |
| --- | --- | --- | --- | --- | --- | --- | --- | --- |
| U-Net | 0.9334 | 0.9354 | 0.8737 | 0.9330 | 0.9034 | 0.8933 | 0.9267 | 0.9141 |
| U-Net /w | 0.9446 | 0.9450 | 0.8778 | 0.9425 | 0.9161 | 0.9102 | 0.9361 | 0.9246 |
| U-Net++ | 0.9401 | 0.9416 | 0.9266 | 0.9367 | 0.9210 | 0.9189 | 0.9346 | 0.9313 |
| U-Net++ /w | 0.9463 | 0.9467 | 0.9340 | 0.9383 | 0.9316 | 0.9265 | 0.9388 | 0.9375 |
| SE U-Net | 0.9275 | 0.9285 | 0.9149 | 0.9298 | 0.9069 | 0.9147 | 0.9277 | 0.9214 |
| SE U-Net /w | 0.9372 | 0.9406 | 0.9219s | 0.9213 | 0.9282 | 0.9269 | 0.9331 | 0.9299 |
| Ux-A4 | 0.9507 | 0.9517 | 0.8167 | 0.9497 | 0.9065 | 0.9015 | 0.9428 | 0.9171 |
| Ux-A4 /s | 0.9562 | 0.9548 | 0.9239 | 0.9553 | 0.9393 | 0.9334 | 0.9523 | 0.9450 |
| Ux-A4 /s/w | **0.9640** | **0.9651** | 0.9291 | **0.9621** | 0.9440 | 0.9320 | 0.9559 | 0.9503 |
| Ux-A10 | 0.9462 | 0.9428 | 0.8625 | 0.9472 | 0.9138 | 0.9135 | 0.9365 | 0.9232 |
| Ux-A10 /s | 0.9594 | 0.9590 | 0.9310 | 0.9550 | 0.9402 | 0.9353 | 0.9507 | 0.9472 |
| Ux-A10 /s/w | 0.9613 | 0.9618 | **0.9514** | 0.9619 | **0.9559** | **0.9500** | **0.9608** | **0.9576** |

**Fig. 5.**
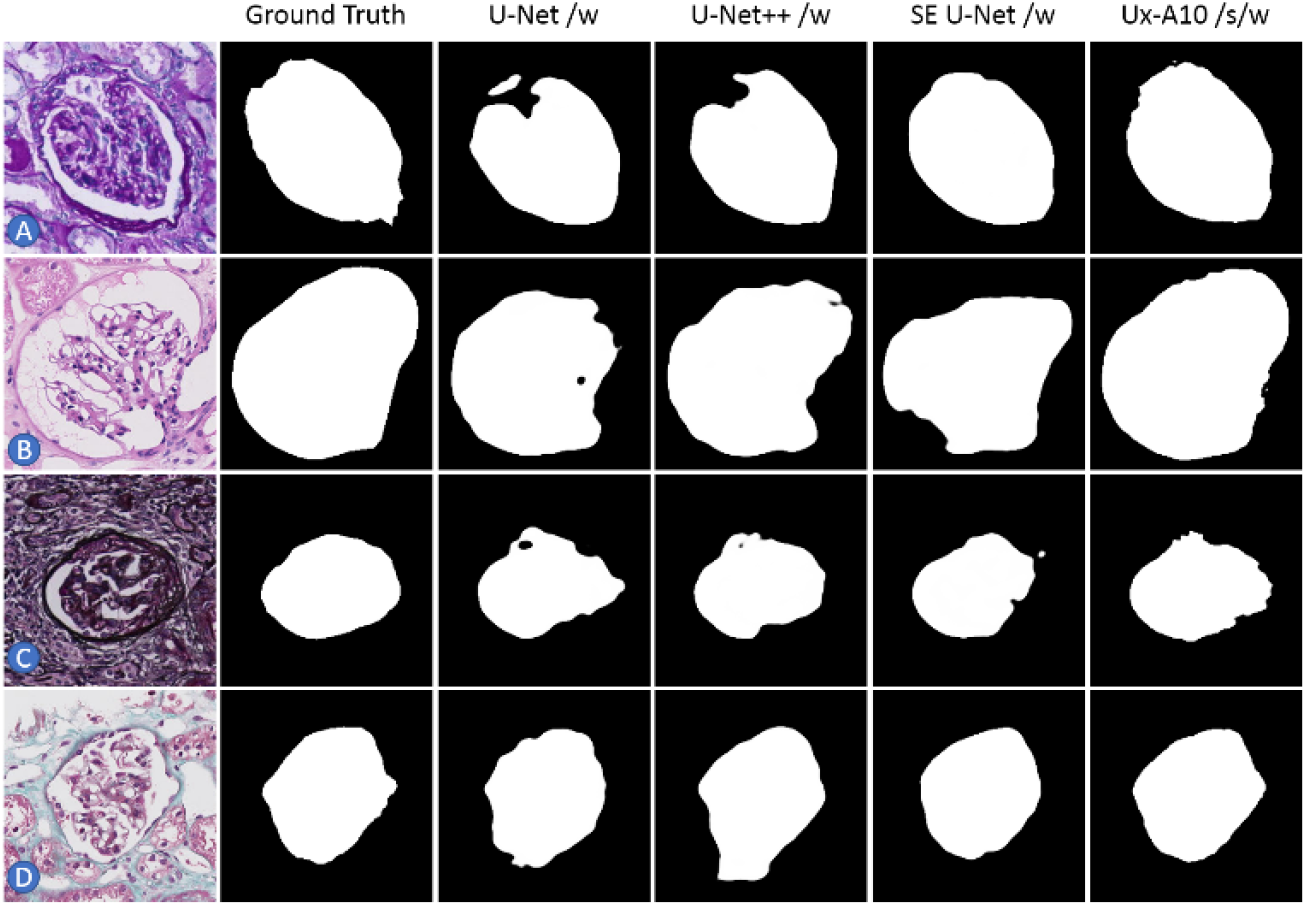
Comparison between U-Net, U-Net++, SE U-Net and proposed framework. A: PAS, B: H&E, C: PASM, D: MT

## 4 Conclusion

We introduced a detection-based framework to segment glomeruli from whole slide imaging of renal biopsy specimens. We explored to use pre-trained classification network to construct and initialize the segmentation network and achieved large performance gain. We proposed a weakly supervised training strategy to resolve the problem of insufficient and inaccurate data annotations. The strategy can also be used to fine-tun the model automatically to adapt more and more new staining even after the algorithm deployed.

## References

1. Gadermayr, M., Appel, V., Klinkhammer, B., Boor, P., Merhof, D.: Which way round? A study on the performance of stain-translation for segmenting arbitrarily dyed histological images. In: MICCAI (2018).

2. Lampert, T., Merveille, O., Schmitz, J., Forestier, G., Feuerhake, F., Wemmert, C.: Strategies for training stain invariant CNNs. arXiv preprint 1810.10338 (2018).

3. Ren, S., He, K., Girshick, R., Sun, J.: Faster R-CNN: Towards real-time object detection with region proposal networks. In: NIPS (2015).

4. Ronneberger, O., Fischer, P., Brox, T.: U-Net: convolutional networks for biomedical image segmentation. In: MICCAI (2015).

5. Reinhard, E., Adhikhmin, M., Gooch, B., Shirley, P.: Color transfer between images. IEEE Computer Graphics and Applications 21(5), 34–41 (2001).

6. Ioffe, S., Szegedy, C.: Batch normalization: accelerating deep network training by reducing internal covariate shift, In: ICML (2015).

7. Simonyan, K., Zisserman, A.: Very deep convolutional networks for large-scale image recognition. In: ICLR (2015).

8. Oquab, M., Bottou, L., Laptev, I., Sivic, J.: Is object localization for free? weakly-supervised learning with convolutional neural networks. In: CVPR (2015).

9. Zhou, B., Khosla, A., Lapedriza, A., Oliva, A., Torrsalba, A.: Learning deep features for discriminative localization. In: CVPR (2016).

10. Yu, F., Wang, D., Shelhamer, E., Darrell, T.: Deep layer aggregation. In: CVPR (2018).

11. Zhou, Z., Siddiquee, M., Tajbakhsh, N., Liang, J.: UNet++: A nested U-Net architecture for medical image segmentation. In: Proc. of Deep Learning in Medical Image Analysis and Multimodal Learning for Clinical Decision Support, 3–11(2018)

12. Roy, A.G., Navab, N., Wachinger, C.: Concurrent Spatial and Channel Squeeze & Excitation in Fully Convolutional Networks. In: MICCAI (2018)

